# Spatial Analysis of T Cell Clonality in Autoimmune Kidney Disease Using TRV Probes

**DOI:** 10.1101/2025.08.29.673064

**Authors:** Cedric Ly, Darius P. Schaub, Robin Khatri, Zeba Sultana, Annika Boxnick, Zheng Song, Tobias Huber, Thorsten Wiech, Eva Tolosa, Ulf Panzer, Stefan Bonn, Christian Krebs, Immo Prinz

**Author notes:** contributed equally (Cedric Ly, Darius P. Schaub, Stefan Bonn, Christian Krebs, Immo Prinz). Correspondence: Immo Prinz.

## Abstract

Hypothesizing that the localization of T cell clones correlates with immune function, our goal was to develop an unbiased method to study the spatial distribution of T cells in tissues. We created an *in situ* hybridization panel with 248 probes that identify immune and tissue cell types, and 132 probes for all variable TRAV, TRBV, TRGV, and TRDV gene segments. Applying this approach to analyze renal biopsies from patients with autoimmune kidney disease, combinations of TRV segments provided spatial information about T cell clonality. Confined clusters of clonally related αβ T cells were found in proximity to increased numbers of antigen-presenting cells, B cells, and other T cells, reflecting local immune cell interactions. γδ T cells were more frequently located outside or at their periphery of T cell infiltration areas. In conclusion, integrating spatial information with TCR clonotype analysis provided new insights into the organization of immune responses at the tissue level.

## Introduction

Clonal expansion is a hallmark of adaptive immune responses^1^, and it is conceivable that expanded T cell clones with the same antigen specificity perform similar immune effector functions in similar locations. Although αβ T cells are primed to differentiate and expand in secondary lymphoid organs, they often exert their effector functions in tissues, such as the kidney^2^. That way, the distribution of naive and expanded central or effector memory T cell subsets across lymphoid and non-lymphoid tissues depends on their differentiation state^3, 4^. Once activated, they can access organs such as the skin, lung, and kidneys^5^ and persist as tissue-resident memory T cells through local homeostasis^6, 7, 8^. Within their target tissue, the clonal distribution of T cells is contingent on cytokine gradients and signals received through direct interaction with tissue and immune cells. At the same time, peptide/MHC-specific triggering of their TCR is likely to induce local proliferation and clonal aggregation in regions with high cognate antigen abundance. Recent studies using spatial VDJ sequencing have demonstrated that T cell clones in tissues are not randomly distributed but are organized in specific spatial patterns that often correlate with their functional roles and the local microenvironment^9, 10, 11, 12, 13^. However, current VDJ sequencing approaches do not reach cellular resolution, and thus, it is difficult to define the cellular interactions of individual T cell clones.

To overcome these limitations, we developed a novel method to investigate the spatial distribution of T cell clones. To this end, we generated an *in situ* hybridization panel comprising probes for all human variable (TRV) TRAV, TRBV, TRGV, and TRDV gene segments, along with sufficient probes for cell-type-specific genes. This allows us to identify immune and tissue cell types and states at single-cell spatial resolution. We established an analysis workflow to identify the expansion and clustering of T cell clones in tissue sections. Here, we applied this workflow to compare the distribution of T cells in biopsies from patients with anti-neutrophil cytoplasmic antibody-associated glomerulonephritis (ANCA-GN) and healthy controls. ANCA-GN is characterized by local inflammation and pathogenic T cell infiltration and thus provides an ideal system for spatial immune analysis^14, 15^. Beyond this initial application, our platform has the potential to study the clonal distribution of infiltrating T cells in any tissue. Thus, it can serve as a simple and fast tool for gaining new insights into the tissue-level organization of immune responses.

## Results

### Spatial profiling of kidney biopsies using a custom TCR gene panel

To resolve clonal T cell expansions by *in situ* hybridization, we developed a targeted gene panel incorporating 132 probes for T cell receptor variable (TRV) genes (**Fig. 1a**), enabling clone detection based on combinations of TRAV/TRBV co-expression. The remainder of the gene panel included 160 probes for genes selected based on prior studies of inflamed kidney tissue, alongside 188 broadly informative probes from established multi-organ panels (**Fig. 1a**). The design of the TRV probes was based on the pmTRIG database^16, 17^, a population-matched reference derived from the 1000 Genomes Project^18^, which catalogs full-length germline alleles across diverse populations. This enabled the selection of variable gene targets with broad population coverage and high specificity, optimized for spatial resolution of T cell clonality (**Extended Data Table 1**). We applied this panel using the Xenium platform (10x Genomics) on formalin-fixed, paraffin-embedded (FFPE) kidney biopsies from three control individuals and ten patients with Anti-neutrophil cytoplasmic antibody (ANCA-)associated glomerulonephritis (GN) (**Fig. 1b**). T cells play a major role in ANCA-GN and localized immune infiltrations are particularly distributed around glomerular regions^14, 15^. After applying a series of quality control steps, including filtering out 25 TRV probes due to a lack of T cell-specific expression, removal of three probes with atypical co-expression patterns, and merging of twelve TRV families comprising cross-reactive probes, we retained 35 TRAV, 36 TRBV, 10 TRGV, and 3 TRDV probes (**Fig. 1c** and **Extended Data Fig. 1a-d**). After quality control filtering, the final probe set permitted a total of 30 TRGV/TRDV and 1,260 TRAV/TRBV combinations (**Extended Data Table 2**), thereby enabling the robust identification of γδ and αβ T cell clones within the spatial tissue context.

**Figure 1.**
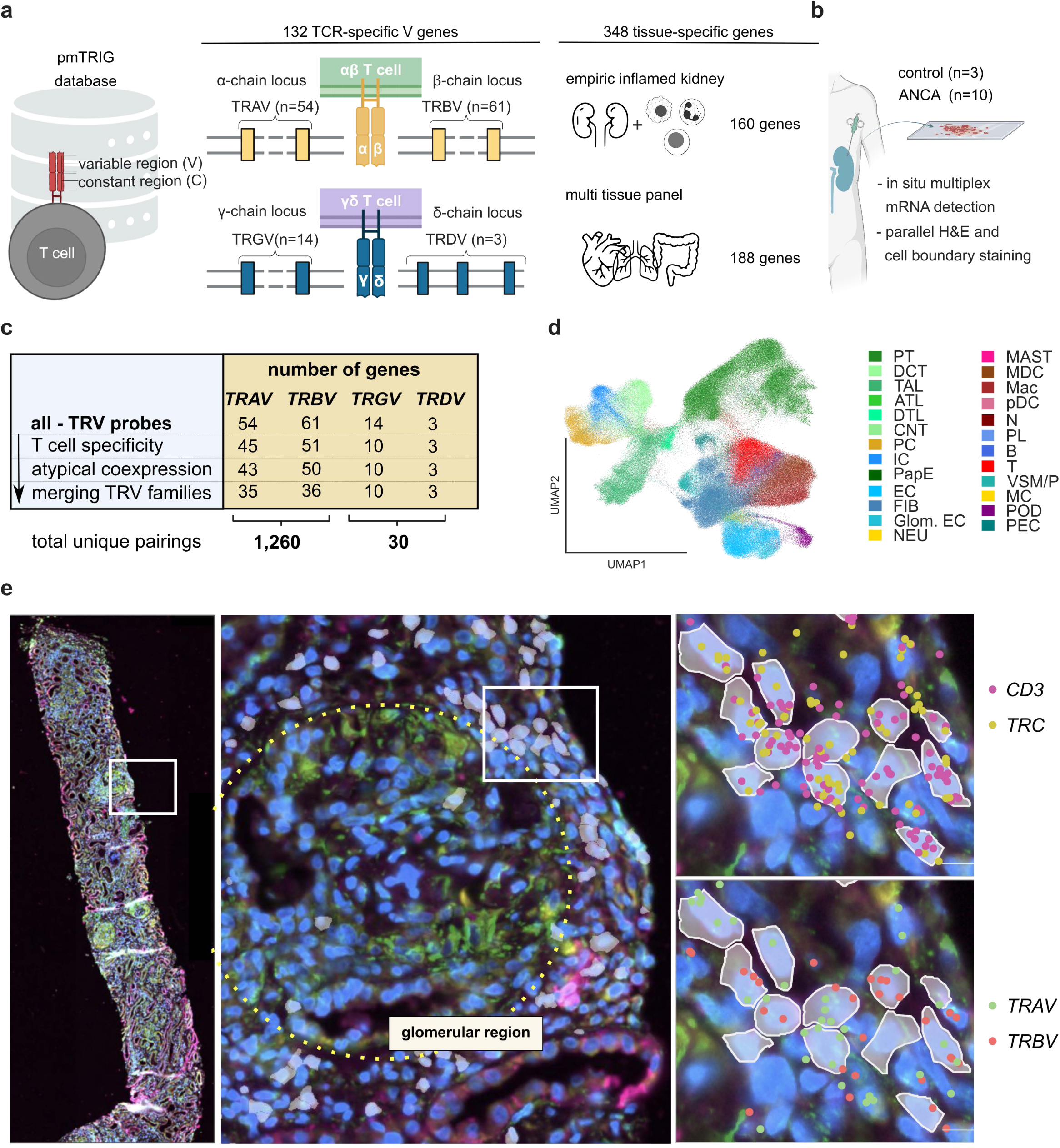
TRV probes reveal the spatial distribution of T cell clones. **a**) Custom gene panel with TRV probes derived from the pmTRIG database, along with probes targeting inflamed kidney tissue and probes from a standard multi-tissue panel. **b**) Application of the panel to human kidney biopsies using high-plex spatial transcriptomics with subcellular resolution. Overview of the TRV gene panel following exclusion of non-specific or cross-reactive probes, retaining sufficient coverage to assess 1,260 potential TRAV/TRBV combinations. Uniform Manifold Approximation and Projection (UMAP) of all cells across the kidney biopsies. T cells are highlighted in red. **e**) Representative ANCA biopsy (left), zoom-in on a glomerular region (middle), and detection of αβ T cells using padlock probes targeting CD3 (CD3E, CD3G, CD3D) and TCR constant regions (TRAC, TRBC) (upper right), corresponding to cells expressing TRAV and TRBV genes (lower right). All results shown are directly derived from the platform’s native output, without additional post-processing.

### TRV gene probe expression maps specifically to T cells

To establish a spatially resolved view of immune cell composition in inflamed kidney tissue, we identified stromal and major immune cell types, including T cells, as shown in the UMAP projection in **Figure 1d**. The cell type classification was performed as described in Sultana et al.^14^. Notably, transcripts specific to T cells, such as CD3E, CD3G, CD3D, TRAC, and TRBC, exhibited robust spatial colocalization with the TRAV and TRBV probes. This highlights the specificity of our customized TRV gene panel for T cells (**Fig. 1e** and **Extended Data Fig. 1e**). As anticipated, in inflamed tissue these transcripts were enriched in areas of immune infiltration, particularly surrounding glomerular regions.

### TRV gene usage delineates αβ and γδ T cells and validates MAIT cell identity

To further resolve the distribution of T cell subsets within the kidney microenvironment, we expanded our analysis to include more detailed annotations of T cell subtypes, including CD4^+^, CD8^+^, regulatory T cells (Tregs), mucosal-associated invariant T (MAIT) cells, natural killer T (NKT)-like cells, and gamma delta (γδ) T cells (**Fig. 2a–b**). Comparison of subtype composition between ANCA and control samples revealed shifts in the proportions of CD4^+^ and CD8^+^ T cells, suggesting disease-associated remodeling of the T cell compartment (**Fig. 2c**). Notably, TRV gene expression patterns aligned with expected lineage specificity as TRAV and TRBV probes were predominantly expressed in αβ T cells, while TRDV and TRGV transcripts localized to γδ T cells. In particular, MAIT cells showed elevated expression of TRAV1-2 (**Fig. 2d**), in agreement with known marker genes for this subset, despite TRAV1-2 not being used as an input for subtype annotation. A representative spatial map highlights this mutually exclusive TRV usage, with an αβ T cell expressing TRAV13-2 and TRBV12, and a γδ T cell expressing TRDV2 and TRGV9 genes within the same tissue region (**Fig. 2e**). These results support the utility of TRV gene probes for both validating and refining T cell subtype identities *in situ*.

**Figure 2.**
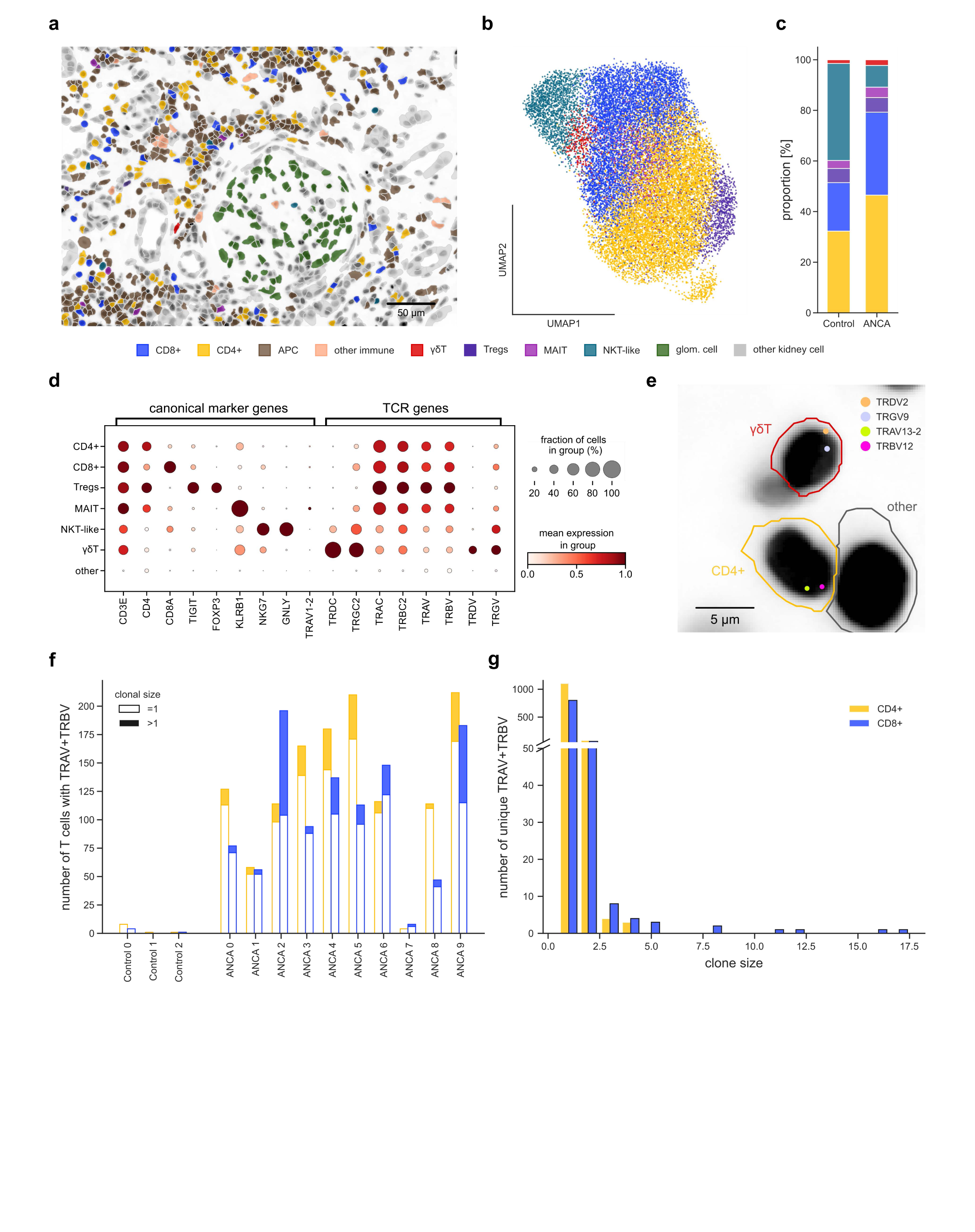
Validation of TRV probes and gene panel performance. **a**) Spatial visualization of a glomerular region from an ANCA biopsy, showing the distribution of kidney-specific cells, immune-related cells, and T cell subsets in color. **b**) Uniform Manifold Approximation and Projection (UMAP) of all T cells and subcluster them by T cell subtype. **c**) Comparing T cell subtype composition between control and ANCA kidney biopsies. **d**) Dot plot showing the expression of subset-defining marker genes and TRV probes across the T cell subsets defined in (b). **e**) Spatial localization of TRAV/TRBV transcripts in an αβ T cell and TRDV/TRGV transcripts in a γδ T cell. **f**) Clonal size (defined by shared TRAV/TRBV combinations) distribution of CD4^+^ (yellow) and CD8^+^ (blue) T cells per biopsy. Single clones are unfilled; expanded clones are stacked and filled. (g) Clonal expansion of CD4^+^ and CD8^+^ T cells across all ANCA biopsies.

### ANCA kidneys comprise more CD4^+^ clones, but CD8^+^ clones show the greatest expansion

To investigate their clonal expansion, we defined T cell clones based on unique TRAV/TRBV combinations. For each sample, we quantified the number of cells sharing identical TRAV/TRBV pairs, allowing us to assess clone sizes within CD4^+^ and CD8^+^ T cell subsets (**Fig. 2f**). Clone size distributions per sample revealed a generally higher number of T cells in most ANCA samples compared to controls, except for sample “ANCA 7” (**Fig. 2f**). Notably, all ten ANCA biopsy samples displayed individual donor-specific repertoires of expanded TRAV/TRBV combinations in CD4^+^ and CD8^+^ T cells (**Extended Data Fig. 2a-c**). Aggregated across all ANCA samples, expanded clones were more numerous in the CD4^+^ compartment, while the largest clone sizes predominantly occurred among CD8^+^ T cells (**Fig. 2g**). Overall, T cell clonal sizes and diversity were consistent with results obtained from re-analysis of our previous single-cell TCR sequencing study of ANCA biopsies^14, 15^, thereby validating our spatially resolved clonal profiling approach (**Extended Data Fig. 3a,b**).

### T cell clonal clusters emerge beyond random spatial aggregation

To understand the spatial organization of expanded T cell clones, we developed a computational workflow that identifies local aggregates of T cells that share the same TRAV/TRBV combination. These were defined as clonal clusters (**Fig. 3a**). Applying this method across all ANCA-GN samples, we detected 145 distinct clonal clusters. To determine whether these clusters arose from random spatial distribution or true biological organization, we performed a permutation test by randomizing clone identities within the tissue while preserving total clone sizes. The observed 145 clonal clusters were significantly more than the expected approximately 100 clusters from permutation analysis (p = 0.001), strengthening the notion of coordinated spatial clustering of T cell clones (**Extended Data Fig. 4a**). Representative examples of clonal clusters are shown in **Figure 3b**, illustrating the localized expansion of specific T cell clones in the inflamed kidney microenvironment.

**Figure 3.**
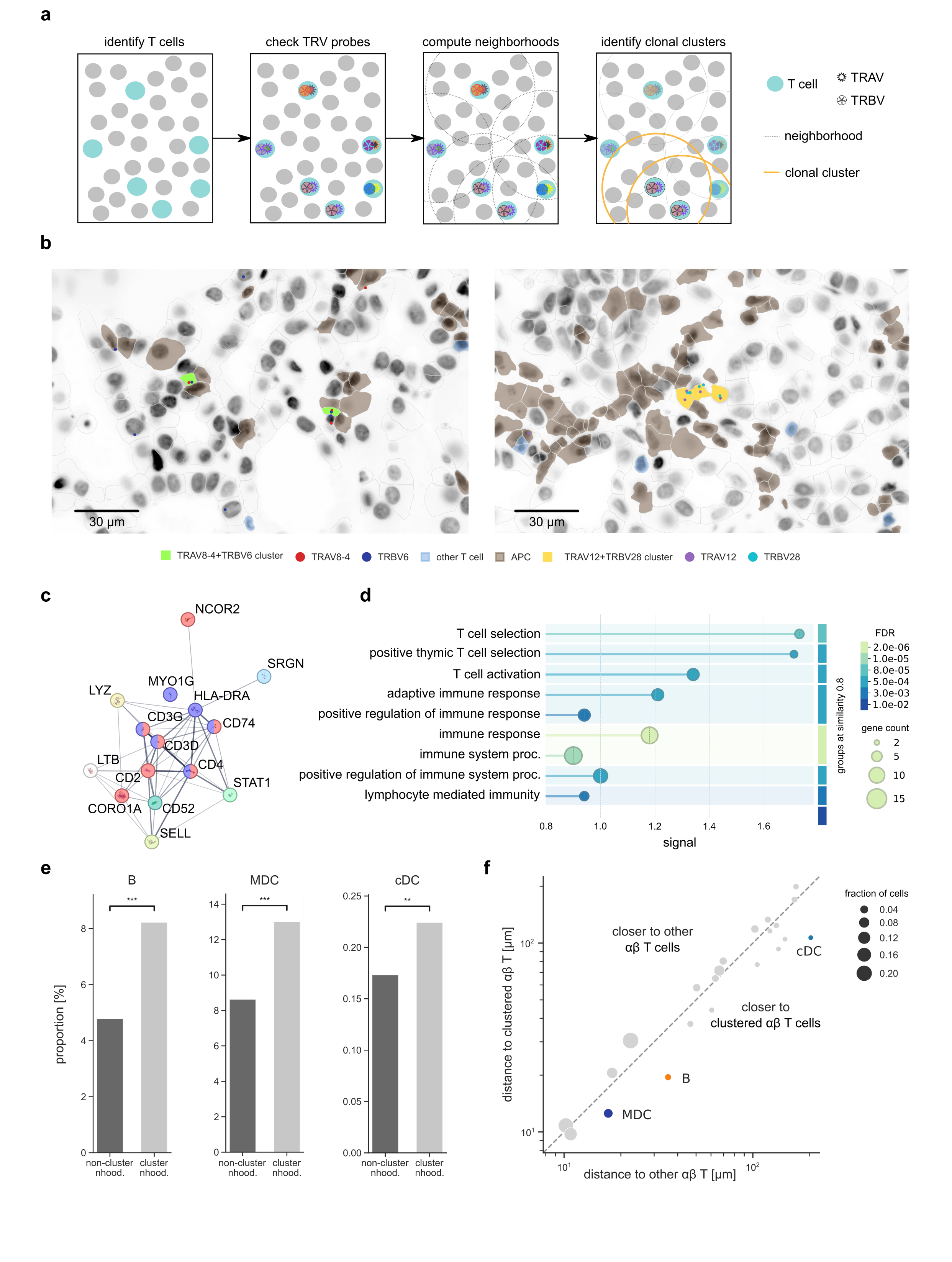
Clonal clusters are activated and localize near antigen-presenting cells. **a)** Schematic workflow to define clusters of T cell clonal clusters and their neighborhood. Various colors of the TRAV and TRBV symbols indicate different TRAV and TRBV genes. Example of clustered T cell clones with highlighted antigen-presenting (APCs) and non-clustered T cells (marked as ‘other T cell’) in the neighborhood. **c,d**) Differential gene expression analysis between clonal cluster T cells and non-clustered T cells (c) shows differentially expressed genes and (d) highlights the pathways enriched among these genes. **e**) APCs are more abundant in the local environment of clonal T cell clusters compared to non-clustered T cells. **f**) Comparison of the average distance from each cell type to clonal clusters versus other T cells. The identity line indicates equal proximity. Points below the identity line indicate shorter distances to clusters.

### Clustered T cell clones exhibit distinct transcriptional signatures

To investigate the molecular profile of T cells within clonal clusters, we performed a differential gene expression analysis comparing them to non-clustered T cells. This analysis identified a distinct set of genes that were upregulated and associated with immune activation and effector function (**Fig. 3c** and **Extended Data Fig. 4b**). Pathway enrichment analysis of these differentially expressed genes (DEGs) revealed overrepresented biological processes related to T cell activation and antigen response, further underscoring their functional distinction (**Fig. 3d**). Leveraging the spatial resolution enabled by inclusion of TRV probes in our gene panel, we analyzed the cellular neighborhoods surrounding clonal clusters versus other T cells. We found that myeloid dendritic cells (MDCs), conventional dendritic cells (cDCs), and B cells, all of which possess antigen-presenting capabilities, were significantly enriched near clonal clusters (**Fig. 3e**). This enrichment was confirmed by shorter distances between APCs and clonal clusters compared to other T cells (**Fig. 3f**), confirming that these antigen-presenting cells are closer to clonal clusters than to non-clustered T cells. This spatial relationship is also visually evident in the representative clonal cluster images (**Fig. 3b**).

### γδ T cells adopt distinct spatial niches from αβ T cells in tissue

Next, to explore the spatial distribution of γδ T cells relative to conventional αβ T cells, we first defined T cell–dense domains within the tissue, which were largely composed of αβ T cells. We then assessed the positioning of γδ T cells in relation to these T cell-infiltrated regions. Interestingly, γδ T cells were more frequently located outside of these high-density T cell areas, often accumulating at their periphery (**Fig. 4a**). This spatial separation was quantified by comparing the distribution of αβ and γδ T cells within and outside the T cell–dense domains, revealing that a significantly lower proportion of γδ T cells were found within these regions (**Fig. 4b**). A permutation test further confirmed that this underrepresentation of γδ T cells in infiltrated areas was unlikely to occur by chance (**Extended Data Fig. 5a**). Spatial neighborhood analysis revealed distinct cellular environments around γδ T cells compared to αβ T cells, with a selective enrichment of neighboring cell types such as other γδ T cells and plasma cells (**Fig. 4c**), suggesting unique tissue interactions for this T cell subset.

**Figure 4.**
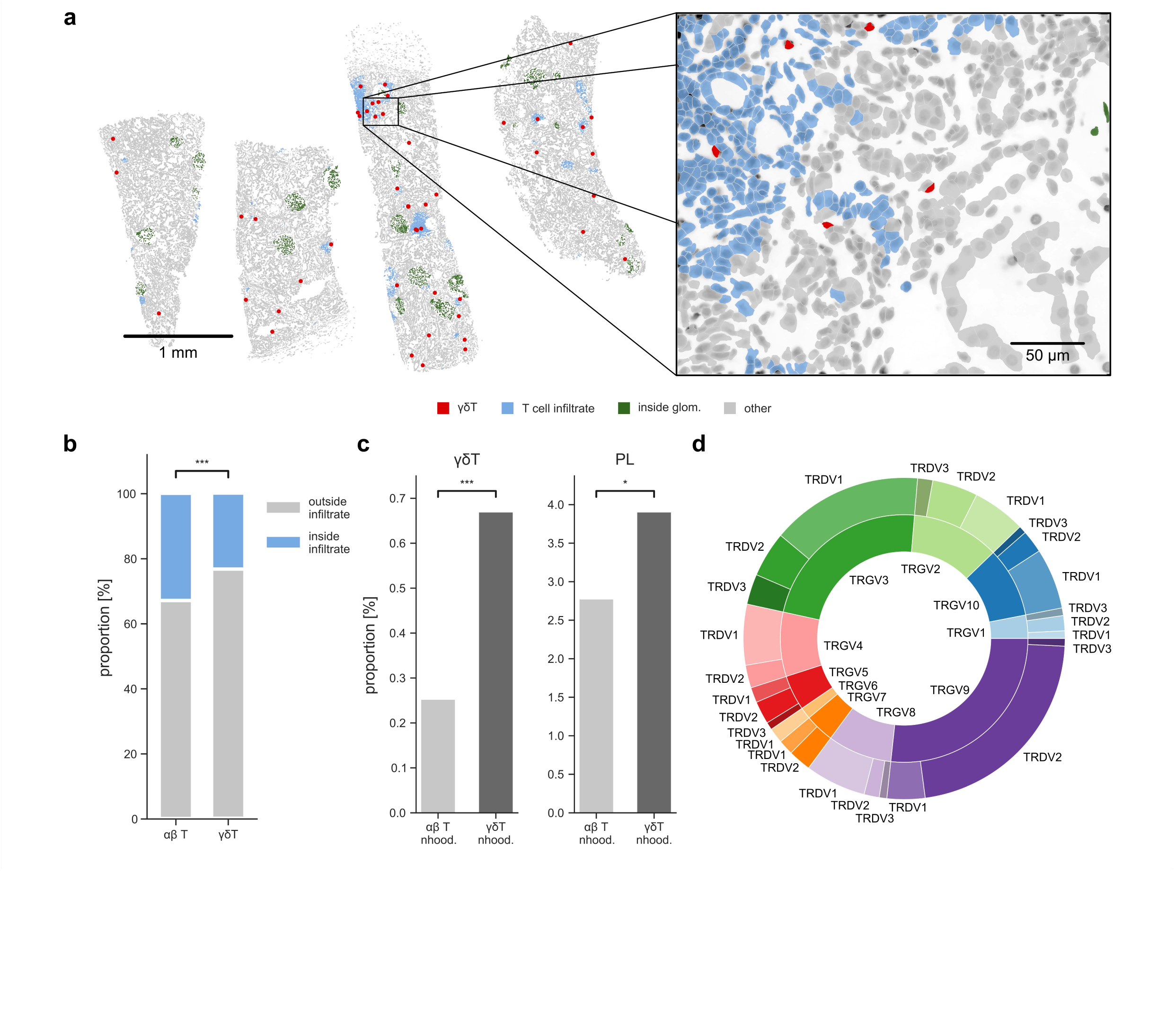
γδ T cells show distinct spatial distributions. **a**) Spatial map from an ANCA biopsy showing T cell–infiltrated areas, glomeruli, and γδ T cells. Magnification highlights γδ T cells located around T cell dense regions. **b**) Comparison of the proportion of γδ versus αβ T cells located within T cell–dense (infiltrated) regions. **c**) Neighborhood cell type composition of γδ T cells compared to αβ T cells, highlighting distinctions in their local cellular environments. **d**) Overview of the γδ T cell repertoire across 10 ANCA kidney biopsies, defined by TRGV/TRDV gene pairings.

In addition, we examined the TRV gene usage across γδ T cells and observed diverse TRDV and TRGV combinations (**Fig. 4d**). The inner circle of the plot reflects the distribution of TRGV genes across all γδ T cells, while the outer ring further subdivides them based on their pairing with one of the three TRDV genes. The expression of the TRDC and TRDV genes is more specific to γδ T cells than the TRGV and TRGC genes^19^. Therefore, using TRDV probes improved our ability to identify γδ T cells, particularly Vγ9Vδ2^+^ cells, compared to classifying them based solely on subclustering. (**Fig. 2b,d** and **Extended Data Fig. 5b-d)**.

## Discussion

This work introduces a novel and simple method for determining the spatial distribution of T cell clones in tissues. Classifying T cell clones using a full panel of TRV *in situ* hybridization probes does not require prior knowledge of the expected CDR3 sequences of the investigated TCRs or their subsequent sequencing. The 1,260 calculated TRAV/TRBV and 30 TRGV/TRDV combinations should provide sufficient information on the spatial distribution of T cell clones in tissues. This assumption was validated by two observations: First, αβ and γδ T cells could be clearly distinguished and displayed different localization patterns in ANCA-GN biopsies. Second, within samples, T cells annotated as CD4^+^ or CD8^+^ rarely exhibited expansions with overlapping TRAV/TRBV combinations. Third, each biopsy sample displayed a unique repertoire of expanded CD4^+^ or CD8^+^ T cells. However, classifying T cell clones based solely on their TRAV and TRBV gene expression is only an approximation because several clones may coincidentally use the same TRAV/TRBV combinations. Another limitation of the study was the imperfect sensitivity and specificity of the TRV panel. However, these results are within the expected range of signal-to-noise ratio for experimental *in situ* hybridization probes. Based on our quality control results, these parameters can be improved for future studies.

In this context, it may be interesting to revisit the issue of allelic exclusion of TRBV and TRAV genes, respectively^20^. While there is stringent allelic exclusion at the TRB locus, approximately 30% of all αβ T cells express two TCR α-chain mRNAs. Conversely, γδ T cells can express TCR β-chain mRNAs, and αβ T cells can express TCR γ-chain mRNAs^19^. Therefore, features of allelic inclusion, including pseudogenes, will increase the number of potential TCR rearrangements and can serve to detect clones with increased specificity using our TRV probe panel.

Since Ig heavy and light chains may be present at much higher abundance per cell, it is tempting to consider designing a similar approach to investigate the spatial distribution of B cell clones. However, it has proven difficult to find primers and probes targeting specific IG V region sequences, while consensus primers are rather easy to find^21^. Additionally, the individual transcriptomes of IGHV genes exhibit significant genetic diversity, largely due to polymorphism and copy number variation^22, 23^. Finally, possible somatic hypermutation would further compromise IGV gene specificity.

While the sensitivity and specificity of TRV probes are sufficient to detect and group individual T cell clones in tissues, this technology competes with other approaches that have their own advantages and disadvantages. One straightforward approach is to use V gene-specific monoclonal antibodies for immunofluorescent staining. However, available reagents only cover a small selection of TRBV and an even smaller selection of TRAV^24^. Alternatively, peptide MHC complexes can be employed to detect T cells that share a common TCR antigen specificity, which requires prior knowledge of antigenic peptides and careful immunofluorescence microscopy setup^25, 26^. Finally, other solutions for spatial VDJ-transcriptomics have been developed for Ig and αβ T cell receptors^9, 10, 11, 12, 13^. However, these approaches require additional PCR and sequencing steps and do not yet reach cellular resolution.

Overall, we offer a straightforward approach and analysis workflow for determining the spatial distribution of T cell clones in formalin-fixed tissue samples. This application can provide insight into local immune responses in infections, cancers, and autoimmune diseases. However, we do not suggest that using this TRV probe panel become a common, stand-alone procedure for mapping T cells in tissues. As multiplexed *in situ* gene expression profiling technologies are quickly evolving^27^, we predict that these TRV probes will be incorporated into larger future panels to routinely provide additional information on local immune responses.

## Methods

### Gene panel design

Several sources were employed for the selection of a total of 480 genes for the 10x Xenium panel. 132 TRV variable genes with three probes each were designed in cooperation with 10x Genomics. 150 genes were selected using the Spapros algorithm^28^ with a single-cell kidney atlas^29^ and ANCA-GN T cell atlas^30^. The remaining genes were added from a multi-tissue panel based on the following criteria: (1) all genes corresponding to heart and kidney tissues, (2) genes annotated for only one cell type and tissue, and (3) the remaining genes were added from the Spapros run only on the multi-tissue panel, excluding the first two lists.

### Preprocessing and quality control

We qualitatively validated the original staining-based cell segmentation by visually comparing the segmentation masks with the staining images across multiple samples and randomly selected tissue slices. To remove doublets, we excluded cells with an area >400 µm², more than 500 transcripts, or expression of over 100 genes. We also removed low-quality cells with fewer than 10 transcripts and excluded all cells located more than 75 µm away from the main biopsy tissue.

### TRV gene filtering

We developed three filtration steps for the TRV transcripts to ensure high-quality measurements, based on three factors: (1) expression level in T cells, (2) non-physiologic co-expression of multiple TRAV or TRBV genes, and (3) the lack of specificity. Separating background noise from meaningful signal is central to transcriptomic analyses, with gene expression serving as the primary readout. Although the background will always be measured, the signal is expected to be higher. The expression of operational TRV probes is expected to be elevated in T cells. It is not surprising that each cell only has a low number of transcripts for a given gene. Therefore, establishing a threshold for the total number of transcripts per cell would significantly reduce the number of cells expressing that particular TRV gene. The initial filter aims to differentiate between "good" and "bad" probes, which would “pass” or “fail” quality control. Only a subset of cells with identified TRV transcripts are T cells. For reliable probes, the proportion of T cells increases when requiring more than one TRV probe per cell. Probes showing a decrease in this proportion after thresholding were deemed not sufficiently T cell specific and excluded from further analysis.

The probes that considered unspecific were predominantly pseudogenes, a finding that was anticipated. Furthermore, we explored the colocalization of TRV genes in the same cell. It was hypothesized that, given the biological rationale that a single T cell should not express more than one unique TRBV due to allelic exclusion, and not more than two unique TRAV genes. Thus, the colocalization of a given TRBV (or TRAV) probe with every other TRBV (or TRAV) probe would indicate a lack of specificity and lead to “fail”. Next, we observed that some TRV genes colocalized only with a limited number of other V genes, but not across the entire TRV gene panel. These were predominantly families of the same gene. In this case, the decision was made to merge these TRV genes, with the aggregate counts serving as the basis for further analyses.

### High-level cell type annotation

A high-level cell type annotation was performed using a logistic regression classifier, trained on an annotated single-cell reference dataset^31^. The intersection of all genes from the single-cell reference dataset and our customized gene panel was subsequently utilized. It should be noted that all T cell receptor (TRV) genes were excluded from the analysis because of their potential to introduce noise during the classification process. The primary rationale for excluding TRV genes was their specific expression in T cell clones, rather than in T cells as a whole.

### T cell subtype annotation

We further identified T cell subtypes by subclustering the cells previously classified as T cells. In particular, we isolated the classified T cells and computed the 100 most highly variable genes, excluding TRV genes, using the *scanpy.pp.highly_variable_genes* function in Scanpy (v1.10.4). Next, we removed any cells that expressed fewer than 5 genes and labeled them as “unknown”. We then performed median normalization, log1p-transformation, PCA, and Leiden clustering using the corresponding Scanpy functions. To check the marker expression per cluster, we restored the full gene expression per cell and computed differentially expressed genes using the *scanpy*.*tl.rank_genes_groups* function in Scanpy. Using this information, we annotated all subclusters. Afterward, we adapted the high-level T cell cluster label to only contain those clusters that were confirmed to be T cells.

### Definition of clones and comparison to TCRseq

We approximate T cell clones by analyzing the expression of TRAV and TRBV genes. After ensuring probe quality (see paragraph "TRV Gene Filtering"), we identified 1,260 possible TRAV/TRBV pairings. A cell annotated as a T cell is only considered part of a clone if it expresses at least one TRAV and one TRBV gene. To define clones, we implemented a majority-rule approach, assigning each cell to a clone based exclusively on its most highly expressed TRAV and TRBV genes. This helps prevent incorrect clone assignments caused by background expression. Once clones are defined, they can be compared to TCR-seq data from the same condition. TCR-seq provides higher resolution of the T cell receptor, as it includes not only the TRV genes but also the CDR3 region. In this study, we compared clonal expansion and associated phenotypes based on TRAV/TRBV pairings between the TCR-seq data and our Xenium dataset. TCR-seq was performed on CD3/CD45 sorted cells from kidneys of ANCA patients (n=6), sc-RNAseq from which have been previously analyzed by Engesser et al.^30^. TCR-seq data for each sample were assembled by the Cell Ranger software (v7.1.0, 10x Genomics) with the command cellranger vdj using the reference genome (refdata-cellranger-vdj-GRCh38-alts-ensembl-7.1.0).

### Clonal cluster identification

To identify spatial clusters of T-cell clones, we first isolated all T cells, previously annotated as αβT cells, that expressed at least one TRAV and TRBV gene. Next, we identified all cells that expressed a certain TRAV/TRBV combination for all possible TRAV/TRBV pairs. These cell sets could overlap between pairs. We ensured that for each cell expressing multiple TRAV or TRBV genes, only those TRAV or TRBV genes were considered that had the highest expression of all the TRAV or TRBV genes, respectively, in that cell. To identify spatial clusters per cell set, we separated the cell sets spatially and applied the DBSCAN algorithm using *sklearn.cluster.DBSCAN*. We imposed that a CD4^+^ and a CD8^+^ T cell could not both be part of the same cluster, to ensure biological validity. We then merged overlapping spatial clusters across cell sets by assigning the overlapping cells to the largest cluster in which they could be found.

### Clonal cluster validation

We validated the clonal clusters statistically by conducting a spatial permutation test. For this, we randomly shuffled the gene expression among all T cells and recomputed the spatial clusters of T cell clones as outlined previously. Note that this keeps the number of TRAV/TRBV clones constant for each permutation. We repeated this procedure 1,000 times and calculated an empirical p-value comparing the observed number of clonal clusters with the simulated null distribution.

### Clonal cluster analyses

We performed differential gene expression analysis comparing the T cells inside the clonal clusters with other T cells using the *scanpy.tl.rank_genes_groups* function from the Scanpy package. We input all the resulting genes (or their corresponding proteins) with a lower adjusted p-value than 0.05 and positive log2 fold change into the STRING v12^32^ web interface to obtain a network plot indicating functional and physical associations between these proteins. We further generated a pathway enrichment plot using the same web interface.

### Neighborhood analysis

We conducted neighborhood analyses comparing clonal T cell clusters with other T cells and γδ T cells with αβ T cells. Specifically, we compared both the cell type composition in these neighborhoods and the gene expression. In both cases, we defined the cell-specific neighborhood as all other cells within a 25-μm distance. To compare the neighborhood cell type composition, we first summed up the number of cells per cell type and neighborhood and then further summed these across all cells belonging to a group of interest (e.g., clonal T cell clusters).

### Spatial proximity analysis

We analyzed the spatial proximity of clustered αβ T cells to other cell types, especially antigen-presenting cells, and compared it to that of the other αβ T cells. For this, we first determined the distance to the nearest cell of every other cell type for all T cells within every biopsy. Then we calculated the median distance to every other cell type for the clustered and other αβ T cells, yielding a distance matrix.

### Glomeruli annotation

We used NichePCA^33^ to perform a general spatial domain annotation and isolated the glomerulus domain based on canonical marker gene expression, such as *PODXL* and *PECAM1*. We further split the domain into separate glomeruli by constructing a spatial graph between cells with a 20-µm radius cutoff and annotating its connected components. To remove noise, we filtered out all connected components containing fewer than 25 cells.

### T cell infiltrate annotation

T cell infiltrates are spatially connected tissue areas containing a high density of T cells. To identify these areas, we first constructed a spatial graph between cells with a 50-µm radius cutoff and then calculated the relative amount of T cells per cellular neighborhood. All cells containing more than 15% T cells in their immediate neighborhood were labeled as part of a T cell infiltrate.

### Additional γδ T analyses

Inspired by Song et al.^19^, we refined the annotation of γδ T cells by leveraging TRV gene expression. We defined scoring metrics for classical αβ T cells and γδ T cells using the *scanpy.tl.score_genes* function from the Scanpy Python package. The αβ T cell score was based on the expression of constant regions of the α and β chains of the T cell receptor, as well as all filtered TRAV and TRBV probes. In contrast, the γδ T cell score was computed using the constant region of the δ chain and TRDV gene expression. We excluded the γ chain from the γδ T cell score due to lack of specificity, as TRGV transcripts are not excised during T cell development and can be expressed in αβ T cells.

## Data availability

The Xenium and TCRseq data will be made available upon publication. The sequences of all TRV probes are provided in Extended Data Table 1.

## Code availability

The source code to process and analyze the Xenium and TCR-seq data is available at https://github.com/imsb-uke/spatial-tcr.

## Acknowledgements

We would like to acknowledge the support of the Single-Cell Sequencing Core Facility at UKE for sample processing and Ian Fiddes at 10x Genomics for help generating the panel of TRV segment-specific probes. This work was supported by funds from the Deutsche Forschungsgemeinschaft (DFG) with grants SFB1194, project number 264599542, to CL, CK, IP, SB, TH, TW, UP; PR727/11-2 and PR727/12-2 to IP; and KR3483/3-1 to CFK. RK was funded by the grant DFG FOR 5068, DS by DFG CRC 1713.

## Author Contribution

I.P., S.B., and C.K. conceptualized the study. C.L. and D.S. conducted the data analysis. A.B. processed samples. R.K., Z.Su., and I.P. designed the gene panel. C.L. and D.S. processed the data. Z.So. helped with the γδ T cell analysis. I.P., U.P., S.B., C.K., C.L., and D.S. designed the figures. C.L., D.S., and I.P. wrote the manuscript. E.T., T.W., T.H., U.P., and C.K. provided patient samples. I.P., U.P., S.B., and C.K. supervised the study. All authors reviewed and approved the final manuscript.

## Supplementary figure legends

**Supplementary Figure 1.**
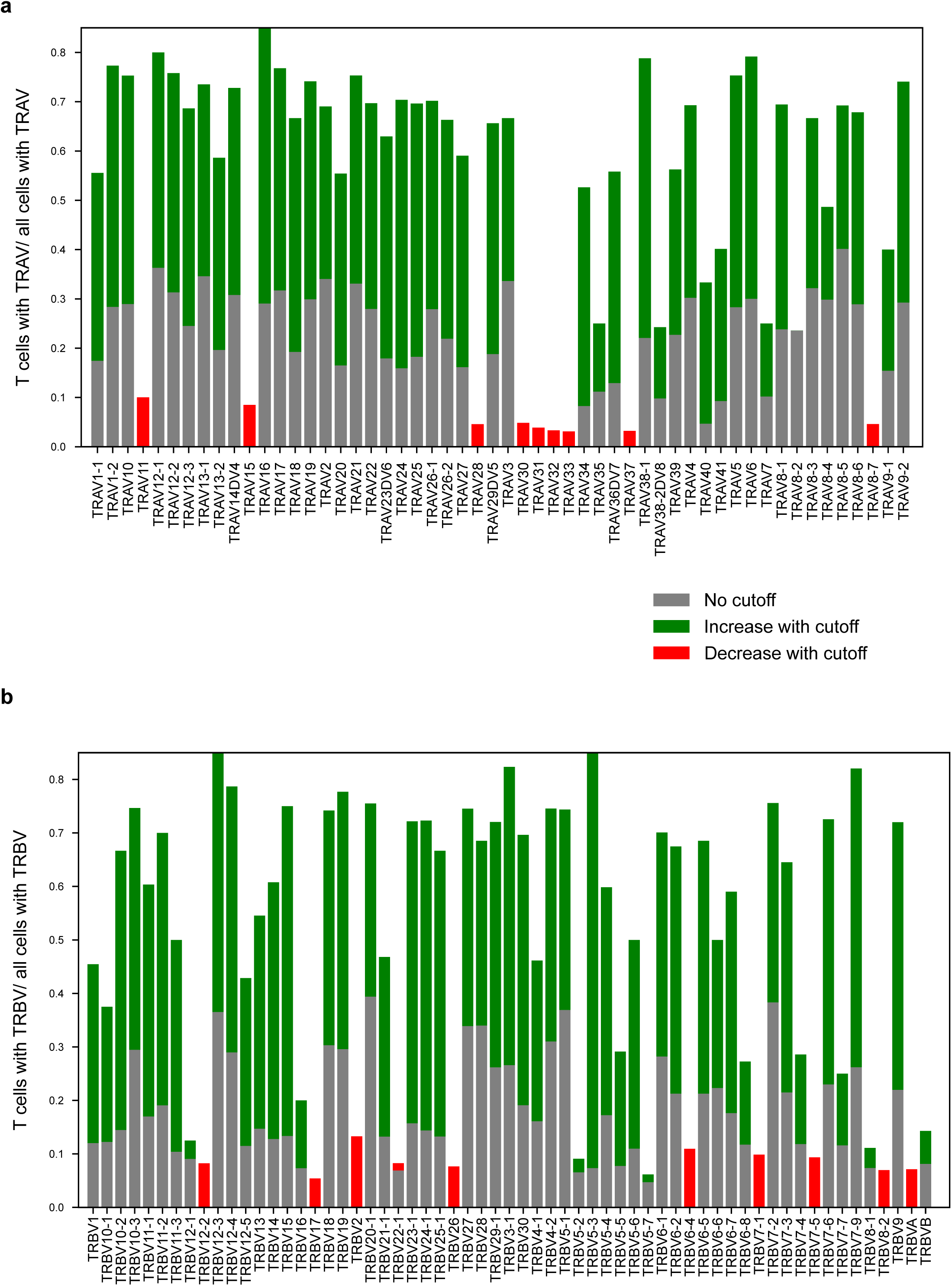
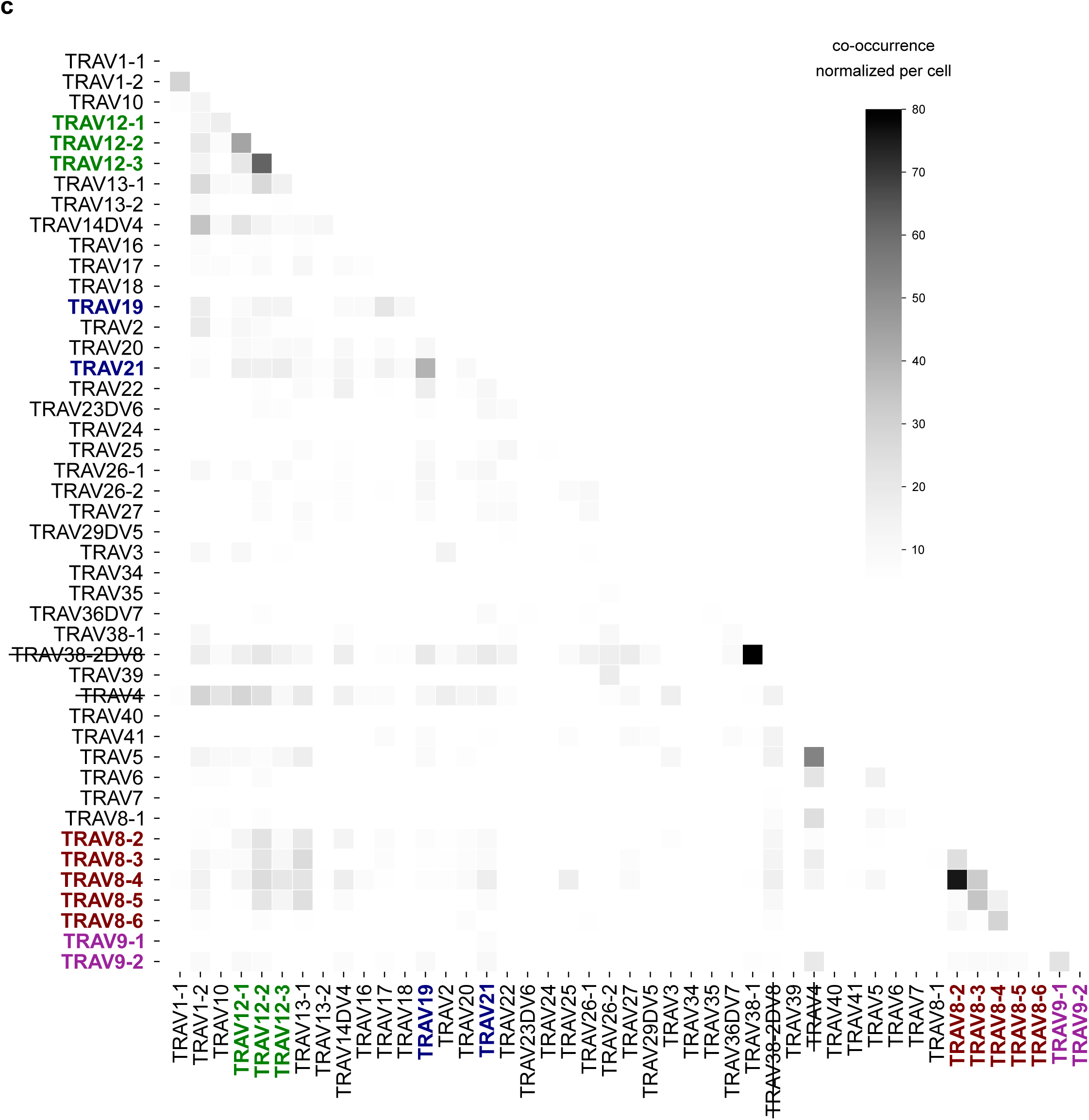
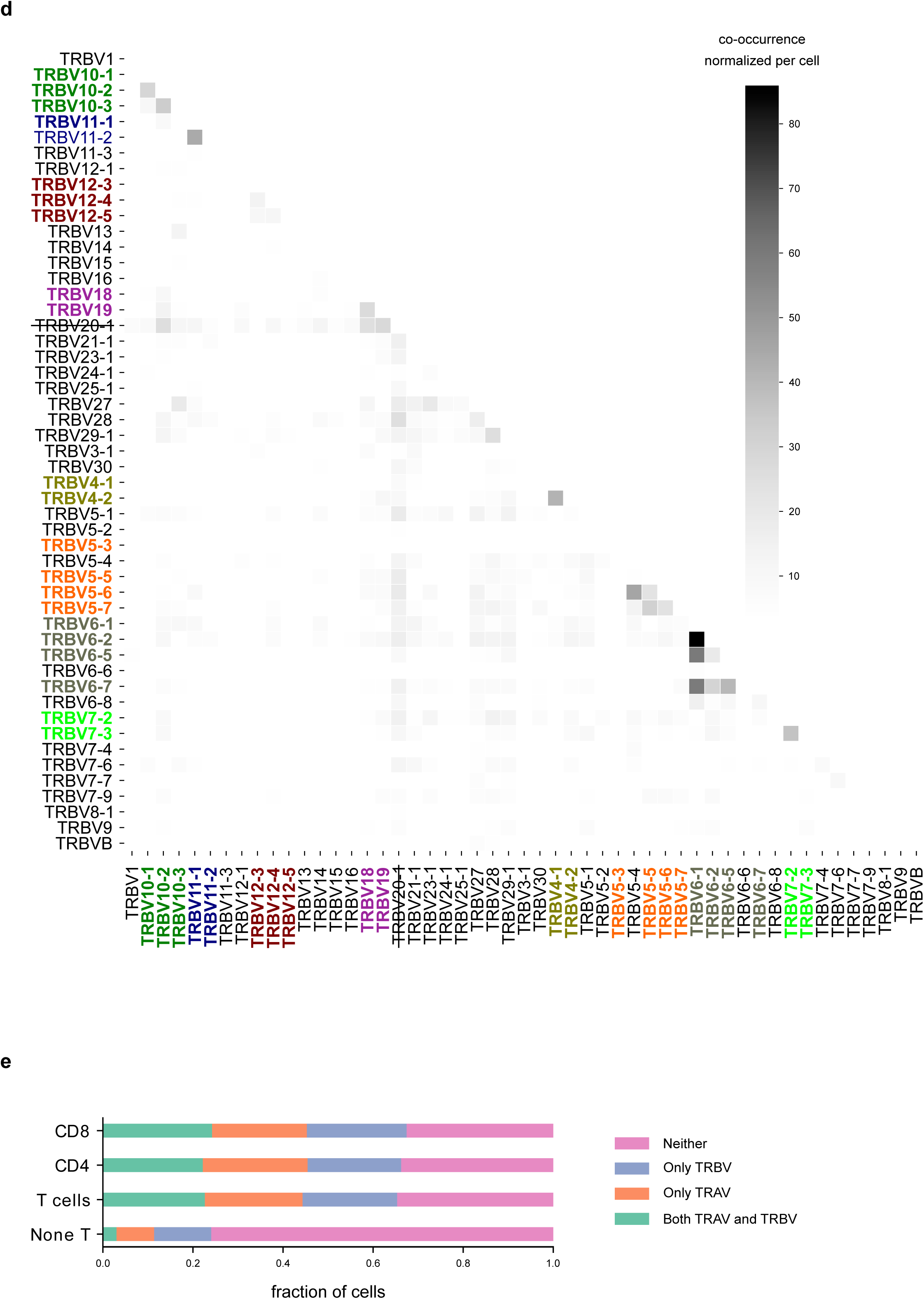
Quality control and filtering pipeline. **a,b**) Bar graphs showing the fraction of T cells among all cells expressing TRAV (a) and TRBV (b). Gray bars indicate the baseline fraction of T cells among all cells where the corresponding TRV probe was detected. Green bars show the increase in T cell fraction when restricting the analysis to cells with at least two detections of the respective probe (indicating a gain). Red bars show the corresponding decrease in T cell fraction under the same threshold (indicating a loss). **c,d**) Co-occurrence heatmaps for TRAV (c) and TRBV (d). TRV genes crossed out are found to be too unspecific, whereas TRV probes in the same color were merged. **e**) Bar graph showing the fraction of cells with detection of both a TRAV and a TRBV probe, only one of the two, or neither.

**Supplementary Figure 2.**
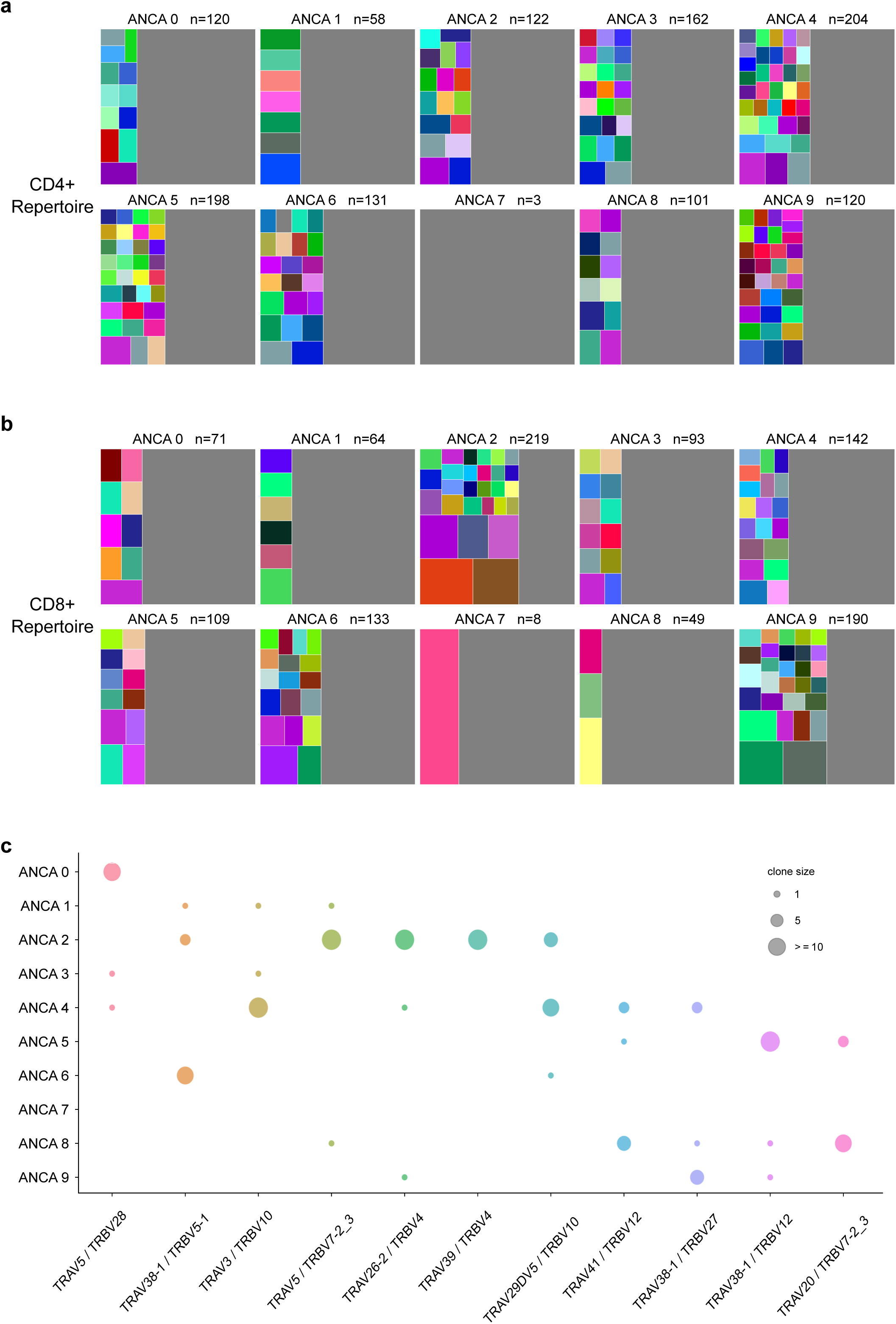
CD4^+^ and CD8^+^ TCR repertoires. **a,b**) Treemap plots display T cell clonal expansions (colored) and singlets (grey) for each ANCA sample. Each rectangle represents a unique TCR (TRAV/TRBV) combination, with area proportional to its frequency within the repertoire. The total repertoire “n” includes all T cells with at least one detected TRAV and TRBV probe. **c**) Scatterplot for various TRAV/TRBV clone sizes across all ANCA samples. The color is clone-specific.

**Supplementary Figure 3.**
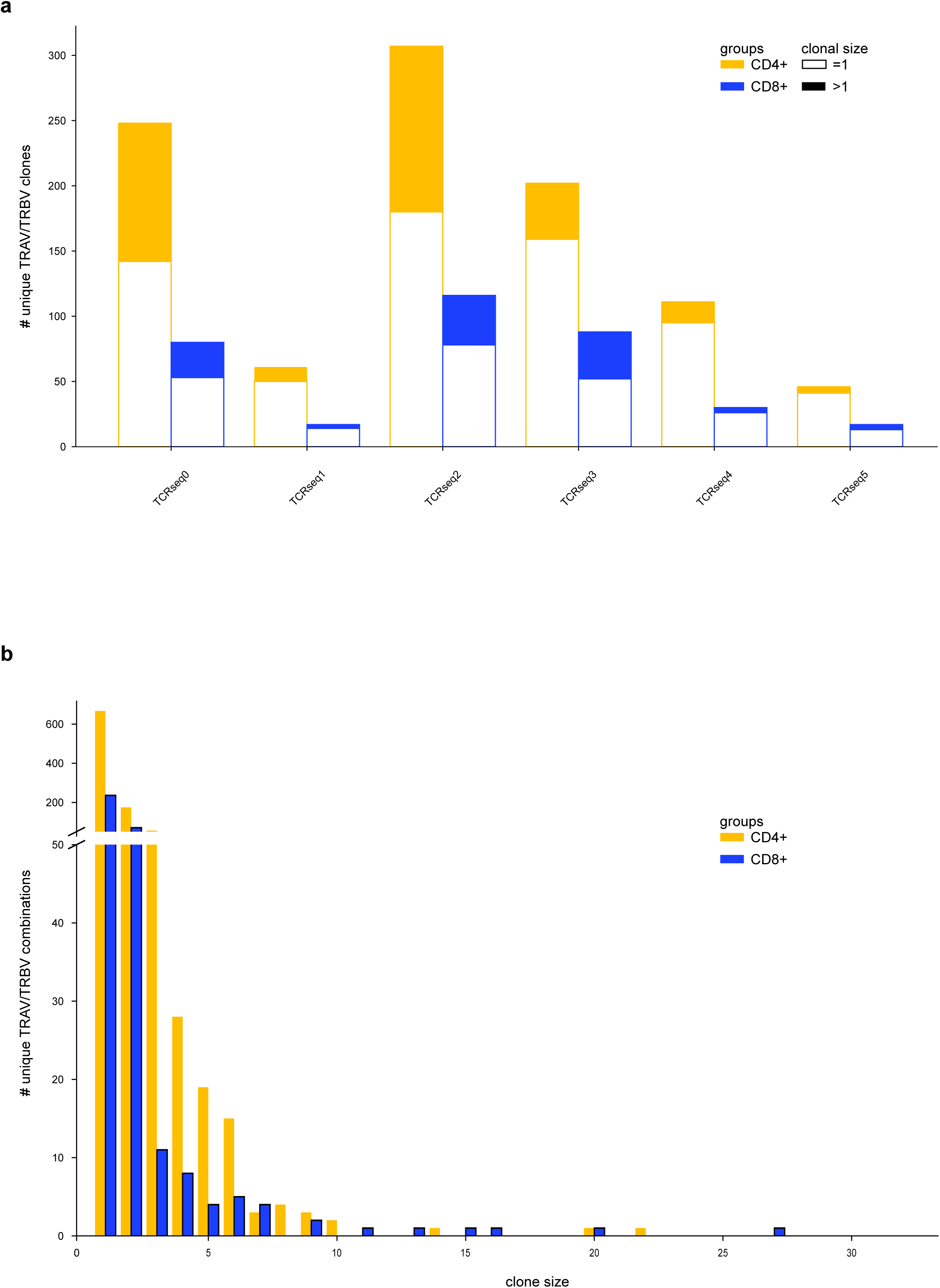
Comparison to single-cell TCRseq data. **a**) Clonal size distribution of CD4⁺ (yellow) and CD8⁺ (blue) T cells in human ANCA kidney biopsies based on shared TRAV/TRBV combinations from referenced TCR-seq data. Singlet clones are shown as unfilled outlines; expanded clones are stacked and filled. **b**) Overall clonal expansion of CD4⁺ and CD8⁺ T cells across all ANCA biopsies.

**Supplementary Figure 4.**
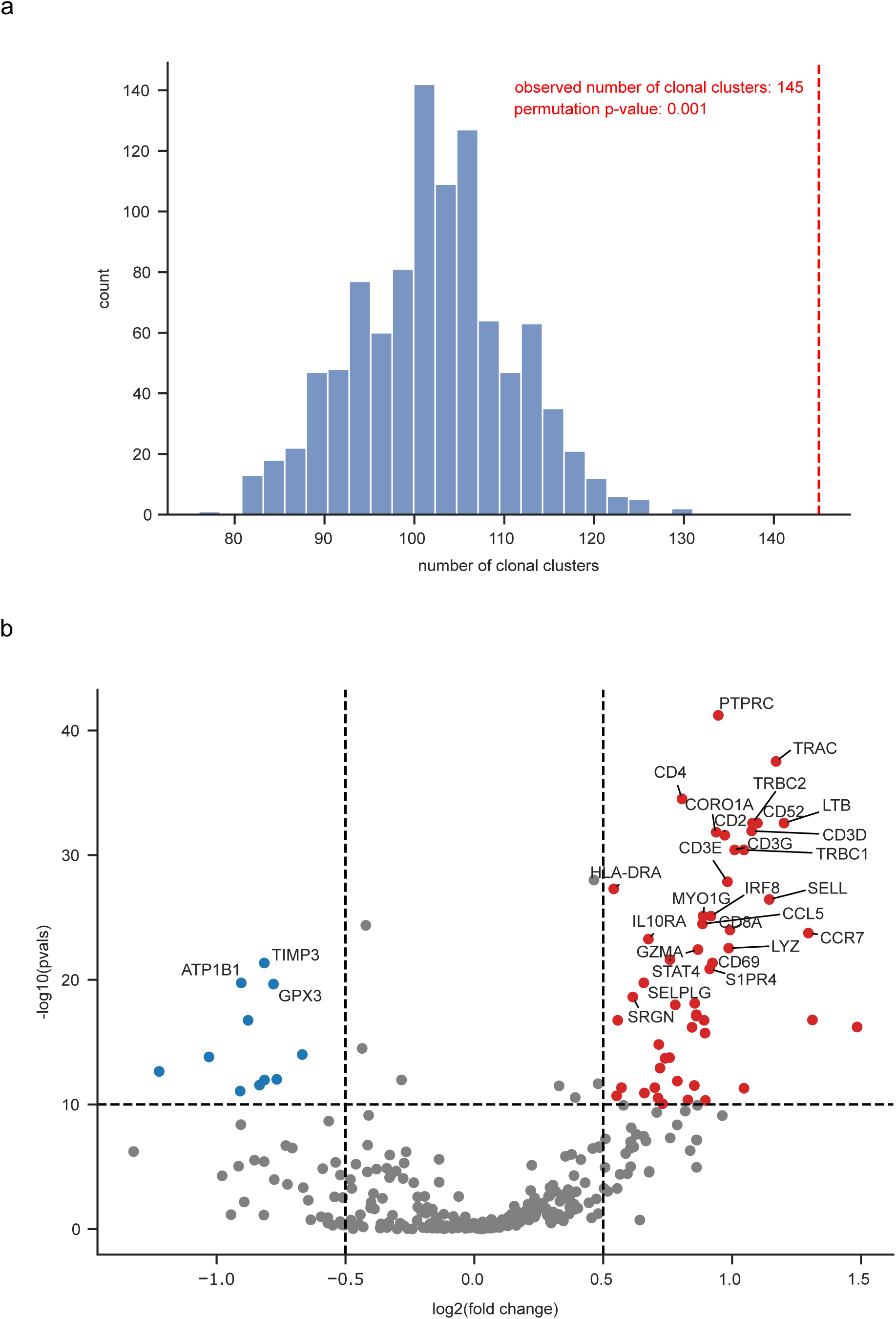
Clonal T cell clusters. **a**) Permutation test illustrating the frequency of found clonal clusters after permutating gene expression 1000 times. The observed number of clonal clusters is marked in red. **b**) Differentially expressed genes visualized as a volcano plot comparing clonal clustered T cells against all other remaining T cells.

**Supplementary Figure 5.**
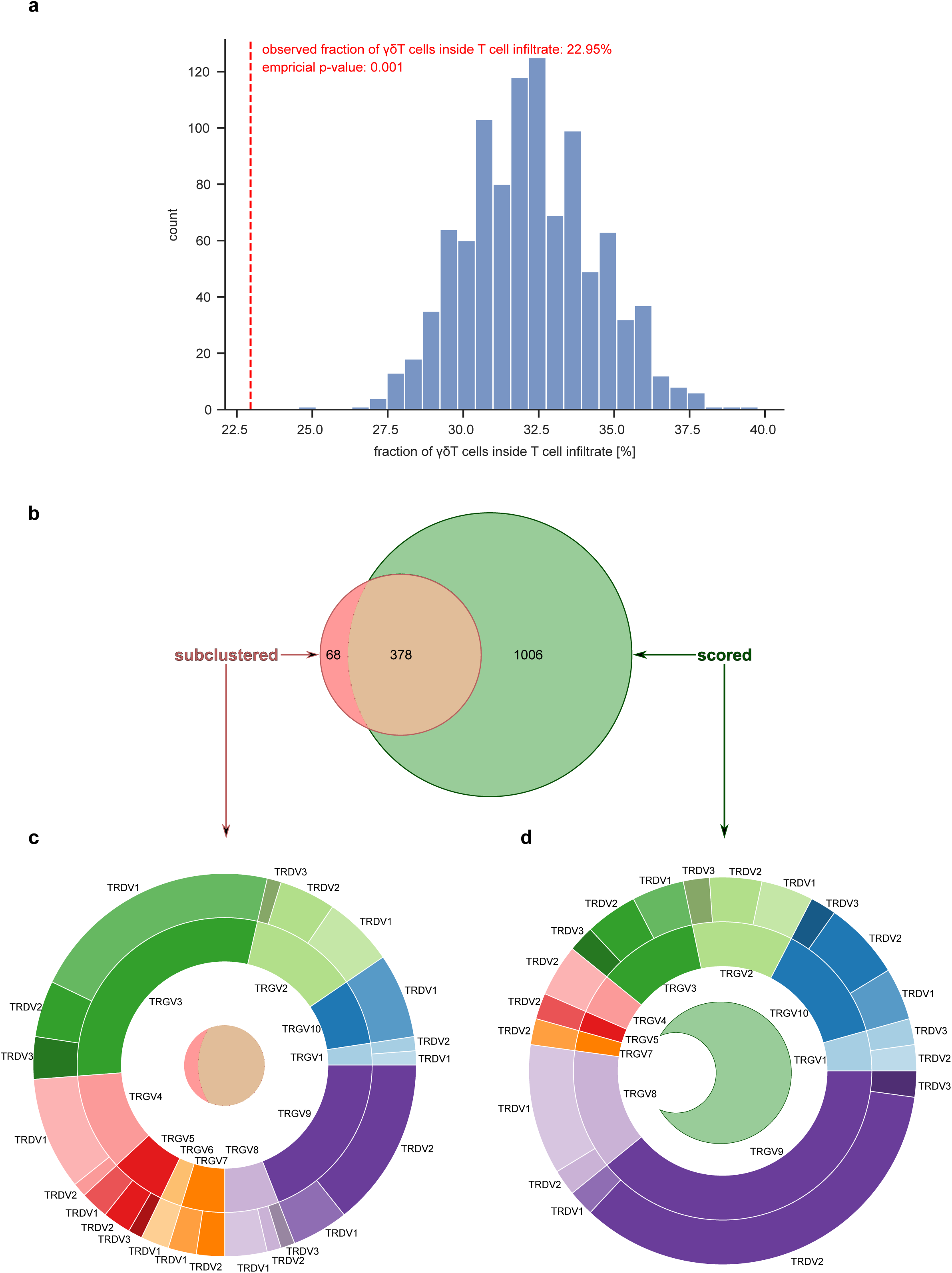
γδ T cell analysis. **a**) Permutation test showing the frequency of observed γδ T cells in infiltrated areas compared to a null distribution generated by random permutations of T cell locations. The observed count is indicated in red. **b**) Venn diagram depicting additional γδ T cells identified by a gene-scoring-based annotation approach. **c,d**) Overview of the γδ T cell repertoire across 10 ANCA kidney biopsies, defined by TRGV/TRDV gene pairings. **c**) Cells annotated via subclustering. **d**) Cells annotated exclusively by the scoring method.

